# ADSC-EVs modulate primary human macrophages to an anti-inflammatory phenotype *in vitro*

**DOI:** 10.1101/2023.05.11.540448

**Authors:** Emma K C Symonds, Bianca Black, Alexander Brown, Ineke Meredith, Margaret Currie, Kathryn E Hally, Kirsty M Danielson

## Abstract

**Background:** EVs released by adipose derived stem cells (ADSCs) have shown promise as a therapeutic for tissue repair and regeneration because of their purported immune-regulatory properties. In this capacity, ADSC-EVs could be beneficial in improving graft retention rates for autologous fat grafting (AFG) post-mastectomy as, currently, grafted tissue rates are reported to be variable and low. Enriching grafted tissue with ADSC-EVs may improve retention rates by modulating macrophages resident within both the breast and lipoaspirate. We aimed to identify key macrophage phenotypes that are modulated by ADSC-EVs *in vitro*.

**Methods:** ADSCs were isolated from lipoaspirates of women undergoing AFG and characterised by flow cytometry and differentiation potential. ADSC-EVs were isolated from cell culture media and characterised by tunable resistive pulse sensing (TRPS), transmission electron microscopy (TEM), and Western blot. Primary monocyte-derived macrophages were polarized to an M1-like (GM-CSF, IFNγ) or M2-like phenotype (M-CSF, IL-4) or maintained (M0-like; M-CSF) and, at the time of polarization, ADSC-EVs were co-cultured with macrophages for 48 hrs. Flow cytometry coupled with high-dimensional analysis was used to cluster macrophages post co-culture. A manual gating strategy was generated to recapitulate these clusters and was applied to a repeat experimental run. Both runs were analysed to examine the prevalence of each cluster, representing a unique macrophage phenotype, with and without ADSC-EVs.

**Results:** Following the addition of ADSC-EVs, M0-like macrophages demonstrated a reciprocal shift of cell distribution from a cluster defined as having a ‘high inflammatory profile’ (CD36^+++^CD206^+++^CD86^+++^; 38.6±14.8% of M1-like macrophages without ADSC-EVs; 16.5±7.0% with ADSC-EVs; p<0.0001) to a cluster with a ‘lower inflammatory’ profile (CD36^+^CD206^+^CD86^+^; 16.6±11.2% to 35±21.5%; p<0.05). There was no shift in M2-like clusters following treatment with ADSC-EVs.

**Conclusions:** ADSC-EVs are complex regulators of macrophage phenotype that can shift macrophages away from a heightened pro-inflammatory state.

## Introduction

Extracellular vesicles (EVs) released by adipose derived stem cells (ADSCs) have shown promise as a therapeutic for tissue repair and regeneration. ADSC-EVs have demonstrated pro-retention and regeneration capabilities, including stimulating wound healing, angiogenesis, and promoting expression of anti-inflammatory mediators (1). As tissue regeneration is highly dependent on the state of the local tissue microenvironment, it is important to study the potential regenerative capabilities of ADSC-EVs in the context of the disease of interest.

Breast cancer is the most prevalent cancer worldwide, with over 2.3 million women diagnosed globally every year (2, 3). Surgical resection is the mainstay of treatment for breast cancer and while the rates of breast cancer are high, the approximate 5-year survival probability following treatment is 90% (4). Post-mastectomy reconstruction has therefore become an integral part of the treatment pathway (5). Autologous fat grafting (AFG), which involves harvesting fat from a donor site and re-injection into the breast wall, is an increasingly desirable breast reconstruction option as it can provide a natural cosmetic outcome, carries low surgical risk and low donor site morbidity, and has a rapid surgical recovery (6). However, whole breast reconstruction with AFG alone is not possible due to the unpredictability of graft survival. Graft retention rates have been reported to be highly variable, ranging from 30-100% (7).

Post resection, and with or without radiation therapy, the breast cavity is a highly inflammatory microenvironment (8, 9) which promotes the infiltration and activation of immune cells involved in wound healing, such as macrophages (10). Successful integration of grafted fat into the breast cavity is therefore likely hindered by this inflammatory microenvironment. The immunomodulatory role of ADSC-EVs on macrophages has previously been highlighted, however, the literature is currently limited to commercially available cell lines or small *in vivo* observational studies. For example, it has been reported that ADSC-exosomes from mice inhibit M1 macrophage polarisation and stimulate M2 macrophage polarisation *in vivo*, reducing the overall inflammatory state of the microenvironment (11). To be a successful therapeutic in the context of AFG, ADSC-EVs need to improve graft retention rates, possibly in part by modulating key immune populations such as macrophages that are resident within the breast cavity at the time of AFG. This study aimed to characterise the effect of ADSC-EVs on primary human macrophages *in vitro* as a step towards understanding how ADSC-EVs could be utilized as a therapeutic for improving AFG retention.

## Materials and Methods

### Recruitment and ADSC Isolation

ADSCs were isolated from the lipoaspirate of 3 women recruited as part of a larger ongoing study on AFG retention. This study has ethical approval from the Health and Disabilities Ethics Committee, New Zealand (19/CEN/23) and all participants provided written informed consent. Fat samples were collected at the time of surgery and transferred to a sterile facility. Cells were isolated using 0.1% Type IA Collagenase (Sigma-Aldrich, MO, USA) and cultured at 37^°^C with 5% CO2. Isolated cells were confirmed as ADSCs by expression of the following phenotype by flow cytometry: CD105^+^CD10^+^CD166^+^CD14^-^CD31^-^ on a BD Biosciences FACS Canto II (Supplementary Figure 1 and supplementary methods). Cultured media (DMEM + 10% EV-depleted FBS + 1% penicillin-streptomycin) was collected at passages 2-5, centrifuged at 2000 x g for 10 mins, and slow frozen to -80°C for downstream EV isolation. To generate EV-depleted FBS, FBS was diluted 1:4 in media, ultracentrifuged at 100 000 x g for 16 hrs, and the supernatant was collected and stored until required.

### EV Isolation and Characterisation

We have submitted all relevant data of our experiments to the EV-TRACK knowledgebase (EV-TRACK ID: EV230010) (12). For EV isolations, cultured media supernatants were thawed at 37^°^C, centrifuged at 2000 x g for 5 mins, and EVs were isolated from 10 mL of supernatant using a combination of size exclusion chromatography (SEC) and ultrafiltration. In brief, EVs were separated using SEC columns (qEV10/35nm) and an automated fraction collector (Izon, New Zealand), fractions 1-4 were collected and pooled, and the total volume was concentrated using Amicon Ultra-15 Centrifugal Filter Units by centrifuging at 4696 x g for 35 mins. For tunable resistive pulse sensing (TRPS) and functional assays, the concentrated product was then diluted as required and analysed fresh. For Western blot analysis, EVs were lysed in Exosome Resuspension Buffer (Thermo Fisher, MA, USA) and frozen at -20^°^C until required. Dummy-EVs (D-EVs) were used as controls for all experiments. D-EVs were defined as media that was cultured in the absence of ADSCs and went through the same isolation process as media cultured in the presence of ADSCs.

The size and concentration of isolated particles was analysed by TRPS using a qNano Gold (Izon). Concentrated particles were centrifuged at 2000 x g for 5 min before analysis on both an NP100 and NP200 nanopore (calibrated to CPC100 and CPC200 beads, respectively). For all samples, the voltage was 0.32 V, stretch was 46.99 mm, and current was 118 nA. All samples were analysed until a particle count of >500 was reached. Results were analysed using Izon Control Suite (version 3.3).

Transmission Electron Microscopy (TEM) was used to visualise EV morphology using 300 mesh formvar/carbon coated copper grids (GSCU300CH-50, ProSciTech, Queensland, Australia) and 4% uranyl acetate stain. Images were analysed on a JEOL JEM2100F microscope using JEOL EDS software (JEOL Ltd, Tokyo, Japan).

Lysed EV samples, as well as controls, were thawed at room temperature prior to being analysed by Western blotting under reducing conditions (using DTT) using CD9, TSG101, Calnexin, and Apolipoprotein B (APOB). ADSC lysates were used as a positive control for CD9, TSG101, and Calnexin, while a visually “milky” plasma sample, (indicating high lipoprotein content), was used as a positive control for APOB (Supplementary Table 1).

### HUVEC Model to Demonstrate ADSC-EV Viability

HUVECs (ATCC, VA, USA) were resuspended in media at 1×10^5^cells/mL and 500 μL was added on top of 300 μL domes of Matrigel (Corning Life Sciences, NY, USA) covering wells of a 24 well plate. ADSC-EVs were added to the media at a concentration of 1×10^5^ particles/mL, based on a previous dose response experiment (Supplementary Figure 2), with an equal volume of D-EV used for control wells. Cells were incubated for 5 hrs before the media was changed to media + 1% Calcein AM stain (Thermo Fisher) and incubated for a further hr. At 6 hrs, the media was replaced, and cells were visualised under a fluorescent microscope (CKX53, Olympus, Tokyo, Japan). Images were analysed using AngioTool (version 6.0) (13) to determine vessel percentage coverage.

### Macrophage Isolation and Polarization

Self-reporting healthy volunteers (n=2) were recruited and provided written, informed consent (HDEC 19/CEN/129). Peripheral Blood Mononuclear Cells (PMBCs) were isolated from the buffy coat of whole blood (20 mL) using RosetteSep human enrichment cocktail (STEMCELL Technologies) and density gradient centrifugation as per manufacturer’s instructions. Isolated cells were cultured in RPMI + 10% FBS + 1% PS at 37°C/5% CO2 for 24 hrs. On day one, media was changed to M0 media (RPMI + 10% FBS + 1% PS + 100 ng/mL M-CSF) and on day seven, media was changed to M1 media (RPMI + 10% FBS + 1% PS + 50 ng/mL GM-CSF + 20 ng/mL IFNγ), or M2 media (RPMI + 10% FBS + 1% PS + 50 ng/mL M-CSF + 10 ng/mL IL-4) or macrophages were maintained in M0 media as above (M0-like phenotype). ADSC-EVs from three consecutive patients were isolated as previously described. 1×10^5^ EVs/mL, diluted in appropriate media, from each patient were added separately to M0-like (M-CSF), M1-like (GM-CSF + IFNγ), and M2-like (M-CSF + IL-4) macrophage cultures of each healthy volunteer at the time of polarization. Equal volumes of D-EVs were used as a control. Macrophages plus ADSC-EVs or D-EVs were then cultured together for 48 hrs.

### Flow Cytometry and High Dimensional Analysis

Macrophages were lifted from plates using Accutase (Sigma-Aldrich) and immediately incubated with 1:1000 Zombie-NIR (BioLegend, CA, USA) for 15 mins at room temperature and Fc blocked (2.5% human Fc block, BioLegend) for a further 10 mins at room temperature. Cells were then diluted 1:1 with an 11-colour antibody cocktail (Supplementary Table 2) and incubated at 4°C for 30 minutes protected from light. Cells were diluted and fixed in 0.67% PBS-buffered PFA (pH 7.4), washed and analysed on a 3-laser (violet-blue-red) Cytek Aurora (Cytek Biosciences Inc., CA, USA). Data were unmixed in SpectroFlo (version 2, Cytek Biosciences Inc.). All samples (24 samples; 8 samples per macrophage polarization state) were cleaned (Supplementary Figure 3; FlowJo, version 10.7.1, BD), and live single cells were imported into OMIQ (Omiq.ai) to build a pipeline for high dimensional data analysis. All samples were arcsinh transformed (cofactor 6000) and were projected onto a two-dimensional space with the UMAP algorithm (neighbours=15, minimum distance=0.4, learning rate=1, epochs=200) (14) using the expression of all fluorescence markers, excluding Zombie-NIR viability dye. The dataset was clustered into phenotypically similar metaclusters using the FlowSOM algorithm (distance metric=Euclidean) (15). The number of FlowSOM metaclusters was iterated until the rarest macrophage population determined by manual gating (pan low expressors) was identified by a single metacluster (metacluster 10). Metaclusters were subsequently colour-overlaid onto the UMAP. Marker expression of each metacluster was examined using overlay histograms, and a description of each metacluster was created. A manual gating strategy was generated to recapitulate all 10 metaclusters (Supplementary Figure 4), each of which represents a unique macrophage phenotype, and was validated in a repeat experimental run (Supplementary Figure 5). Within this repeated experimental run, all 10 unique macrophage phenotypes were identified using this FlowSOM-guided manual gating strategy independently by two authors (E.S., K.H.) and shared high concordance. Both experimental runs were analysed using this manual gating strategy (analysed byy E.S.) and examined for the prevalence of each unique macrophage phenotype with and without treatment with ADSC-EVs.

### Statistical Analysis

Results are expressed as mean ± standard deviation, unless stated otherwise. All statistical analysis was performed using GraphPad Prism software version 8. Clusters + D-EVs or ADSC-EVs were analysed using unpaired t-tests and individual marker data were analysed with a Wilcoxin signed rank test. A *p* value of 0.05 or less was considered statistically significant.

## Results

### EV Characterisation

To confirm successful isolation of EVs, size and concentration of the released particles was measured using TRPS. ADSCs released 3.0×10^10^ ± 0.80 and 6.9×10^8^ ± 0.89 EVs/mL of media with a diameter of 118 ± 4 and 269 ± 11 nm using an NP100 and NP200, respectively (Figure 1A-B). As expected, ADSC-EVs expressed TSG101 and CD9 and were negative for calnexin and APOB (Figure 1C). ADSC-EVs were visualised using TEM, with both close up and wide field images taken. Particles demonstrated typical rounded morphology and were observed within the expected size range (Figure 1D). HUVECs were seeded in Matrigel with or without the addition of ADSC-EVs to determine the functional impact of ADSC-EVs on HUVEC tube formation. Relative to D-EVs, HUVECs with ADSC-EVs had an average fold change of 1.28 ± 0.05 in vessel percentage area (*p*=0.0494; Figure 1E-F).

**Figure 1:**
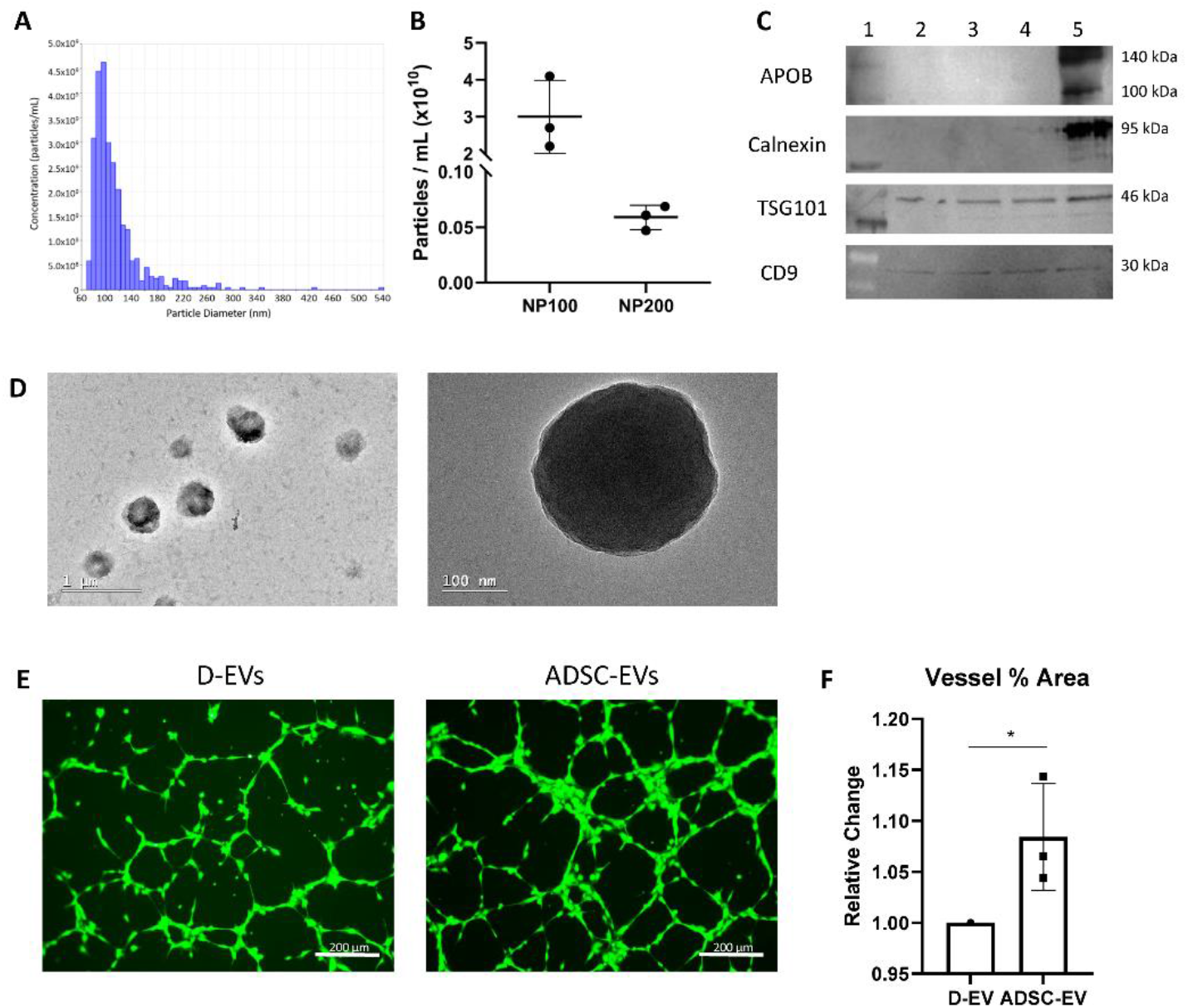
ADSC-EV Characterisation. A) Representative histogram of concentration (particles/mL) versus diameter (nm) following TRPS analysis. B) Concentration (particles/mL) analysed by TRPS comparing an NP100 and NP200 nanopore for three consecutive patients. C) Presence of EV proteins TSG101 and CD9 in lysed EVs in three patient samples (lanes 2-4) and positive control (lane 5). APOB and Calnexin are absent in EV samples, but present in positive control. Ladder is in lane 1. D) Representative TEM images of EVs isolated from patient samples demonstrating typical rounded morphology. E) Representative images of HUVECs seeded on Matrigel with D-EVs (left) and ADSC-EVs (right). F) Percentage of total area taken up by HUVECs when treated with ADSC-EVs relative to D-EVs. Images of three fields of view were taken per patient (n=3) and averaged. Data were analysed using an unpaired t test. \**p*=0.0494.

### Confirmation of Macrophage Polarization

To investigate the role of ADSC-EVs on macrophage phenotype, monocyte-derived macrophages (MDMs) were differentiated in the presence of M-CSF and subsequently polarized into M1-like (GM-CSF + IFNγ) or M2a-like (designated throughout as M2-like; M-CSF + IL-4) and compared to unpolarized M0-like macrophages (maintained with M-CSF). ADSC-EVs isolated from the lipoaspirate of three consecutive patients undergoing AFG surgery were co-cultured with M0-like, M1-like and M2-like macrophages from two self-reporting healthy volunteers. Immunostaining of macrophages with an 11-colour macrophage-centric panel confirmed the successful polarization of M1-like and M2-like macrophages (Figure 2A). The intensity of all fluorescence markers were overlaid onto the UMAP rendering and, based on marker expression, gating on the generated UMAP parameters identified all three polarization states (Figure 2A). The remaining gated region represented macrophages from all three polarization states. Relative to the other polarization states, M1-like macrophages characteristically expressed the highest levels of CD86, CD80, and HLA-DR and had a classical egg-shaped rounded morphology (Figure 2B). M2-like macrophages were the highest expressors of CD206, expressed high levels of CD36 and low levels of CD14 relative to M1-like macrophages, and appeared spindly and stretched. M0-like macrophages were distinct from their M1-like and M2-like counterparts, with high expression of CD64, CD14, and CD163, and had a morphology consisting of a mixture of M1-like and M2-like properties (Figure 2B).

**Figure 2:**
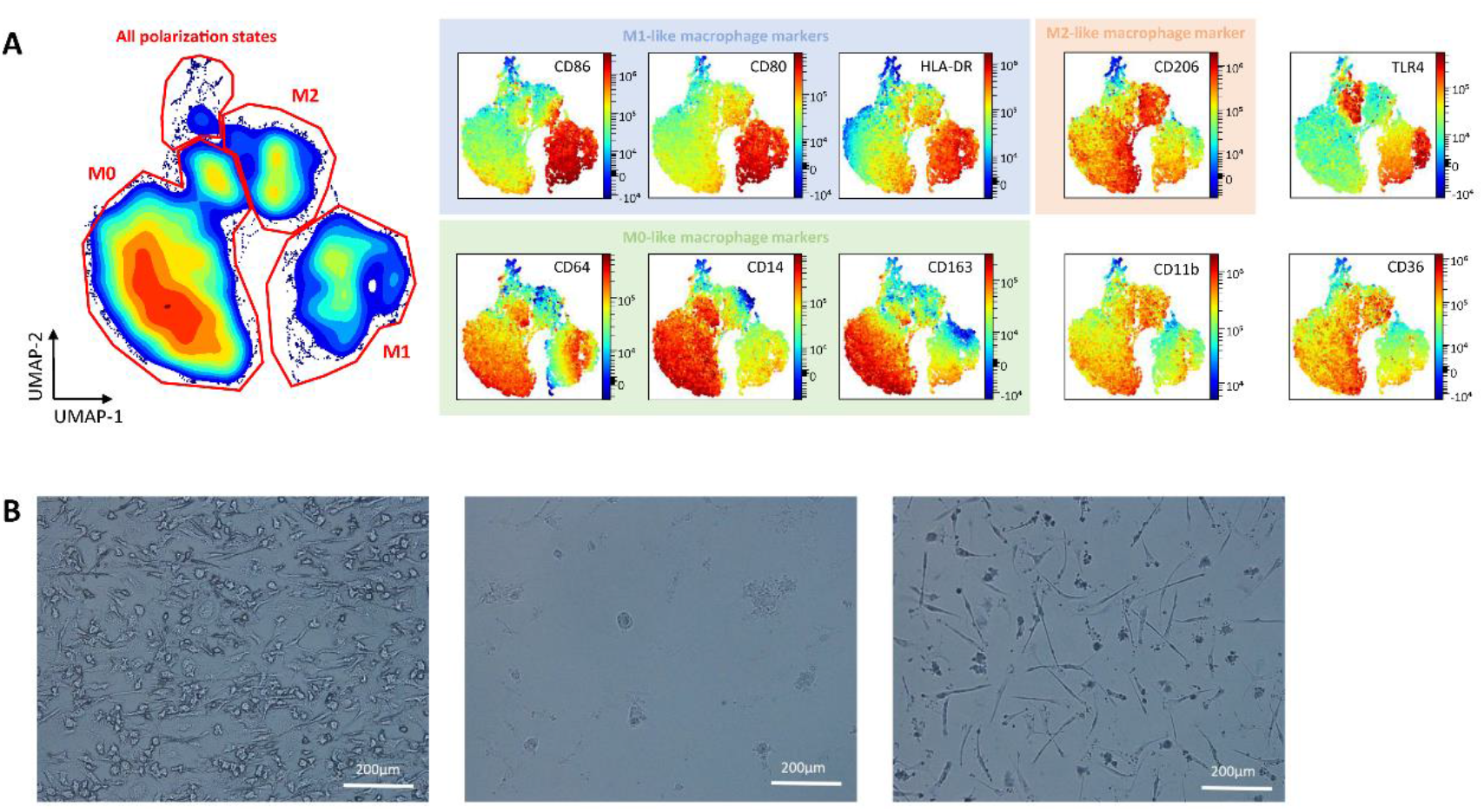
Confirmation of Macrophage Polarisation. A) Immunostaining with an 11-colour macrophage-centric panel and visualization with the UMAP algorithm identifies distinct phenotypes associated with M1-like (GM-CSF + IFNγ) macrophages (CD86++CD80+HLA-DR++), M2-like (M-CSF + IL-4) macrophages (CD206+) and M0-like (M-CSF) macrophages (CD14++). Pan low expressing macrophages were present in all polarization states. B) Representative images of cultured, polarised (left: M0, middle: M1, right: M2) macrophages visualised at 20x on an Olympus BX51 microscope and imaged using an Olympus DP20 camera.

### Flow cytometry and high dimensional analysis identifies unique macrophage phenotypes

FlowSOM clustering identified 10 metaclusters in the dataset, a majority of which were exclusively assigned to one polarization state (Figure 3). One metacluster contained macrophages from all three polarization states (metacluster 10) and another was shared across both M0-like and M2-like macrophages (metacluster 6). Overlay histograms (Figure 3B) were used to tag each metacluster based on their expression pattern and these cluster descriptions (Figure 3C) were used to recapitulate each of these unique macrophage phenotypes by manual gating (Supplementary Figure 4). There was high concordance between manual and unsupervised identification of each metacluster, now defined as unique macrophage phenotypes, as demonstrated by the overlay of manually gated populations onto the UMAP rendering (Figure 3D) and visual comparison with FlowSOM clustering (Figure 3A).

**Figure 3:**
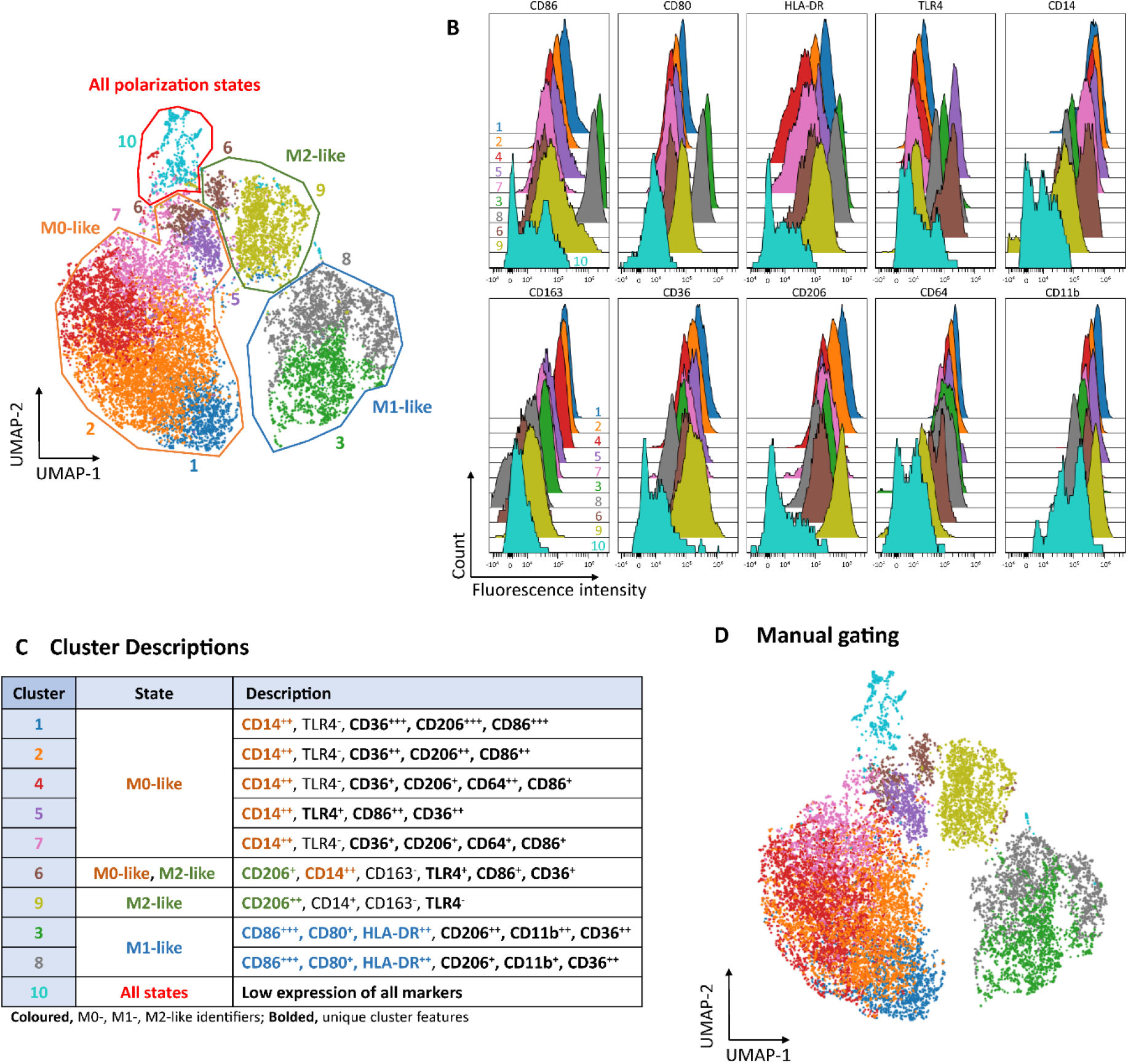
Flow cytometry and high dimensional analysis identifies unique macrophage phenotypes. A) Data was clustered into 10 phenotypically homogenous clusters using the FlowSOM algorithm and each metacluster was colour-overlaid onto the UMAP. B) Marker expression of each metacluster was represented as overlay histograms. C) The overlay histograms were use to tag each metacluster based on their expression pattern. D) Cluster descriptions were used to recapitulate each metacluster by manual gating, and each manually gated population was overlaid onto the UMAP. There was high concordance between unsupervised (A) and manual (D) identification of these clusters.

**Figure 4:**
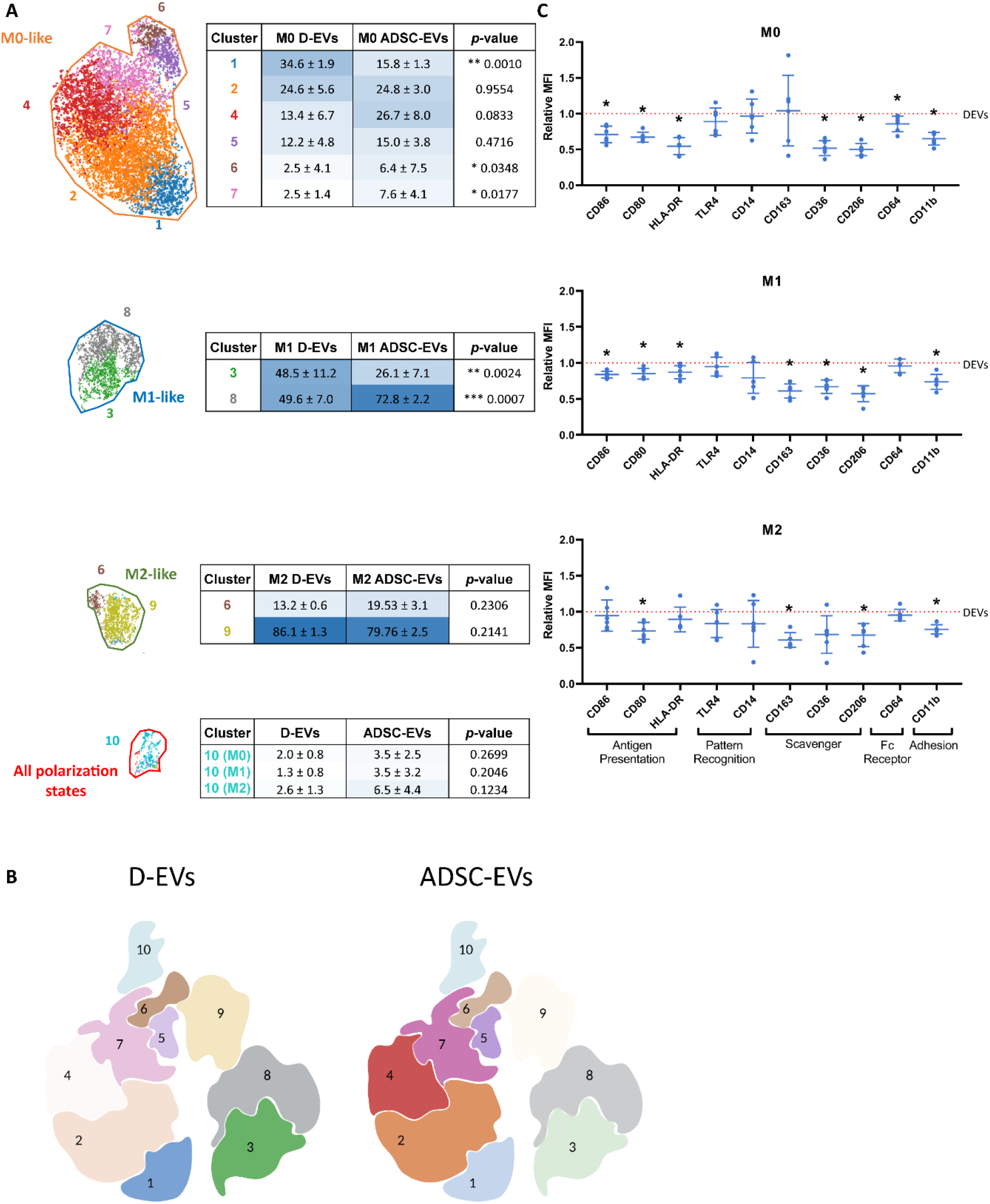
ADSC-EVs modulate macrophage phenotype. A) FlowSOM-guided manual gating was used to assess the effect of ADSC-EV co-culture. ADSC-EVs modulate the proportion of macrophages identified within each metacluster. Values are expressed as mean percentage ± SD. B) Visual representation of the redistribution of macrophages across different metaclusters when treated with ADSC-EVs compared to D-EVs. The darker the colour, the more macrophages represented in the metacluster. C) ADSC-EVs reduce the expression of most markers individually relative to D-EVs (red line). Data were analysed using unpaired t tests. *p*<0.05 was deemed statistically significant.

### ADSC-EVs modulate macrophage phenotypes towards a lower inflammatory profile

The manual gating strategy generated to recapitulate the findings from FlowSOM analysis was validated in a repeat experimental run. All 10 populations were identified in the repeat run (Supplementary Figure 5), and the proportion of each population was shown to be relatively consistent across both experimental runs. These runs were combined and analysed for the prevalence of each unique macrophage phenotype with and without treatment with ADSC-EVs. Results are demonstrated numerically in the tables in Figure 4A and visually in Figure 4B.

For CD14^++^ M0-like macrophages, we demonstrate a significant reduction in the percentage of macrophages within Cluster 1, defined as high expressors of CD36, CD206 and CD86 (CD36^+++^CD206^+++^CD86^+++^), with the addition of ADSC-EVs (34.6±1.9% to 15.8±1.3%, *p*=0.0010). ADSC-EVs seem to facilitate a shift of macrophages from high to low expressors of these markers, as we see a reciprocal increase in the percentage of macrophages assigned collectively to Clusters 4 and 7 (both defined as CD36^+^CD206^+^CD86^+^ and separated by their expression of CD64) post-ADSC-EV treatment (16.6±11.2% to 35±21.5%, *p*<0.05). We also demonstrate a significant shift towards enrichment in cluster 6 (CD206^+^CD14^++^CD163^-^ CD86^+^CD26^+^) for M0-like macrophages following ADSC-EV treatment (2.5±4.1% to 6.4±7.5%, *p*=0.0348).

A similar movement towards dampened marker expression is seen in CD86^+++^CD80^+^HLA-DR^++^ M1-like macrophages. Cluster 3 (defined additionally as CD206^++^CD11b^++^CD36^++^CD163^++^) and Cluster 8 (CD206^+^CD11b^+^CD36^+^CD163^+^) were equally represented in macrophages treated with D-EVs (Cluster 3, 48.5±11.2%; Metacluster 8, 49.6±7.0%) but macrophages moved from metacluster 3 (reduced to 26.1±7.1%, *p*=0.0024) to populate metacluster 8 (increased to 72.8±2.2%, *p*=0.0007) post-ADSC-EV treatment. ADSC-EVs did not significantly impact either of the M2-like metaclusters (6 and 9). In all three macrophage polarization states, a small proportion of macrophages were defined as pan low expressors (Cluster 10). When pooled across polarization states, the proportion of these low expressors represent 2.0±1.0% with D-EV treatment and increased to 4.5±3.7% with ADSC-EV treatment (*p*=0.0119).

On an individual marker level, ADSC-EVs reduced the expression of most of the above-mentioned markers for all three macrophage polarization states (Figure 4C). This was found to be consistent when applied to a repeat experiment (Supplementary Figure 6).

## Discussion

We have successfully defined a population of small ADSC-EVs that significantly affects macrophage phenotype across polarization states. EVs were characterised by TRPS, Western blot, and TEM, and have the expected biological activity in a HUVEC tube formation model. FlowSOM-based clustering of monocyte-derived macrophages co-cultured with D-EVs or ADSC-EVs identified 10 distinct metaclusters. Each metacluster, representing a unique macrophage phenotype, was characterised by different expression levels of 10 macrophage-centric markers, including classical M0-like (CD64, CD14, CD163), M1-like (CD80, CD86, HLA-DR), and M2-like markers (CD206) and markers involved in antigen presentation (CD86, CD80, HLA-DR), adhesion (CD11b), pattern recognition (TLR4, CD14), as well as Fc receptor (CD64) and scavenger receptor expression (CD36, CD163, CD206).

All metaclusters were recapitulated by manual gating, and two independent experimental runs were analysed using this gating strategy. Six metaclusters were identified within CD14^++^ M0-like (M-CSF) macrophages and, with ADSC-EV co-culture, there was a significant depletion of macrophages within metacluster 1 (CD36^+++^CD206^+++^CD86^+++^) coupled with significant enrichment within metaclusters 4/7 (CD36^+^CD206^+^CD86^+^) demonstrating a distinctive movement from high to low expression of these functional markers. Within CD86^+++^CD80^+^HLA-DR^++^ M1-like (GM-CSF + IFNγ) macrophages, ADSC-EV co-culture drove depletion of macrophages from cluster 3 (CD206^++^CD11b^++^CD36^++^) in favour of enrichment within cluster 8 (CD206^+^CD11b^+^CD36^+^), again signifying a shift towards ‘low expressor’ macrophages. Interestingly, unlike M0-like and M1-like macrophages, ADSC-EVs did not significantly impact either of the M2-like metaclusters (6 and 9). Macrophages that were defined as ‘pan low expressors’ (metacluster 10) were present across all three polarization states, and significantly enriched in ADSC-EV co-cultures. We conclude that ADSC-EVs have an immunomodulatory role in this context, by inducing a shift in macrophage phenotype from high to low expressors.

Unsupervised clustering and dimensionality reduction using FlowSOM, a self-organizing map algorithm (15), is a powerful tool for identifying and comparing the presence of phenotypically distinct clusters across experimental conditions. In this study, 10 metaclusters successfully captured the rarest macrophage population (‘pan low expressors’) within a single metacluster (metacluster 10), and a FlowSOM-guided manual gating strategy was generated and validated in a repeat experimental run. FlowSOM-guided manual gating successfully identified important, but nuanced, shifts in macrophage phenotype with ADSC-EV co-culture in a manner that is not easily or reproducibly achieved by traditional manual gating alone. Here, this strategy allowed us to identify a global shift towards reduced expression of markers involved in scavenging, adhesion and antigen presentation (16, 17, 18).

The shift to a lower expression profile of these markers involved in these functional processes is promising in the context of graft retention, as increased activation of inflammatory cells in the breast cavity is thought to be one of the contributing factors to low retention rates (19). Macrophages, which are known for their functional role in wound healing and inflammation, are reported to be one of the dominant cell types present in the breast cavity post-surgical resection (20, 21). Traditionally, M2-like macrophages have been investigated in the context of graft retention in the breast cavity, and are known to be associated with increased angiogenesis, increased stem cell recruitment, and overall increased graft weight three months post-surgery in a limited number of animal studies (20, 21). Given the well-characterized immunosuppressive function of M2-like macrophages, the results presented here support the notion that dampening the inflammatory microenvironment within the breast cavity is likely to support increased graft retention.

The growing consensus is that macrophages exhibit a spectrum of polarity, rather than distinct polarizations states (22, 23), such that the breast cavity at the time of AFG is likely to be populated with macrophages of diverse phenotype and function. The immune compartment within normal breast (intraepithelial) tissue is diverse and for the purpose of host defence and immune surveillance. Our best estimates of the composition of the breast immune compartment of cancer survivors undergoing breast reconstruction comes from literature investigating normal breast biopsies (ipsilateral non-adjacent or contralateral from prophylactic mastectomy) at time of surgical resection. Within this literature, there is agreement that (CD68^+^) macrophages (24) are a prominent immune cell, accompanied by significant numbers of T cells (25), Natural Killer cells (26), and dendritic cells (27). In women undergoing oncologic breast reconstruction, the extent of surgical manipulation during resection of the primary cancer, subsequent wound healing, and (possible) radiation therapy is likely to change the immune landscape towards a pro-inflammatory environment (9, 28).

The immune compartment within subcutaneous white adipose tissue is relatively well-established, and highly dependent on the presence of increased adiposity and metabolic dysregulation. Within lean adipose tissue, an anti-inflammatory environment predominates due to the abundance of M2-like (anti-inflammatory) macrophages, accompanied by eosinophils, regulatory T cells (29) and Type 2 innate lymphoid cells. In obesity and T2DM, adipose tissue often displays hallmark shifts towards a pro-inflammatory state including a favouring of M1-like macrophages (over M2-like macrophages) (30), a favouring of Th1 and Th17 CD4+ T helper cells (over regulatory T cells), and an increase in the frequency and activation of CD8+ cytotoxic T cells (31). This pro-inflammatory shift may contribute to reduced graft retention rates in women with obesity and/or metabolic disorders where the introduction of a pro-inflammatory tissue (autologous lipoaspirate) into the breast cavity may tip the balance in favour of adipocyte necrosis and tissue destruction. Here, we demonstrate that ADSC-EVs may facilitate a re-balancing towards an anti-inflammatory environment by modifying macrophage cell phenotype and function, an action which may contribute to successful grafting.

Moreover, successfully utilizing allogenic ADSC-EVs is promising for the potential use of these particles as an “off-the-shelf” therapy, and supports the preferred development of allogenic EV-based therapeutics over autologous transfer of whole-cell ADSC therapies. This study also demonstrated the abundance of EVs isolated from one fat sample (for example, EV-enriched media can be collected from ADSC passages 2-5), as well as a relatively low dose required to facilitate an effect. This highlights the potential for ADSCs as an easily obtainable population of mesenchymal stem cells for therapeutic EV harvest. Finally, we have chosen an EV isolation method that does not utilise ultracentrifugation and is potentially more scalable for therapeutic production platforms. Ongoing work will need to address issues including sterility and good manufacturing process for further development of ADSC-EVs as a therapeutic.

Following this, it would be prudent to investigate the role of ADSC-EVs on other cells types abundant in the breast cavity at the time of AFG, such as fibroblasts, endothelial cells, T cells, neutrophils, and adipocytes (32, 33). While the effects of ADSC-EVs on macrophages described herein are of great value in the wider context of AFG retention, inflammation is not governed solely by macrophages and is one of many processes hypothesised to hinder graft retention rates. For example, angiogenesis and fibrosis are both important processes that contribute to the success of an integrated graft, indicating a need to investigate the potential functional role of ADSC-EVs on other cell types (34, 35). This work has wider implications for EV-derived therapeutics in a number of contexts where macrophages play a role in wound healing. Macrophages are a key regulator of inflammation and are also known to be involved in a vast range of diseases and conditions, such as cancer, chronic inflammatory diseases, and cardiovascular disease (36, 37, 38). The techniques from this study are thus applicable to other disease models, both to identify important but potentially nuanced shifts in macrophage phenotype, as well as assessing the role of ADSC-EVs as potential therapeutic agents.

Here, we demonstrate that ADSC-EVs have an immunomodulatory role on macrophages *in vitro*. This demonstrates potential for the use of ADSC-EVs as a therapeutic in the context of AFG retention.

## Acknowledgements

This study was supported by the Health Research Council New Zealand, Breast Cancer Cure, Breast Cancer Foundation New Zealand and the Marsden Fund.

## Patient consent

Patients and participants have given written, informed consent.

