## Supplementary for "ADSC-EVs modulate primary human macrophages to an anti-inflammatory phenotype *in vitro*"

**Appendix / Supplementary Figures and Methods**

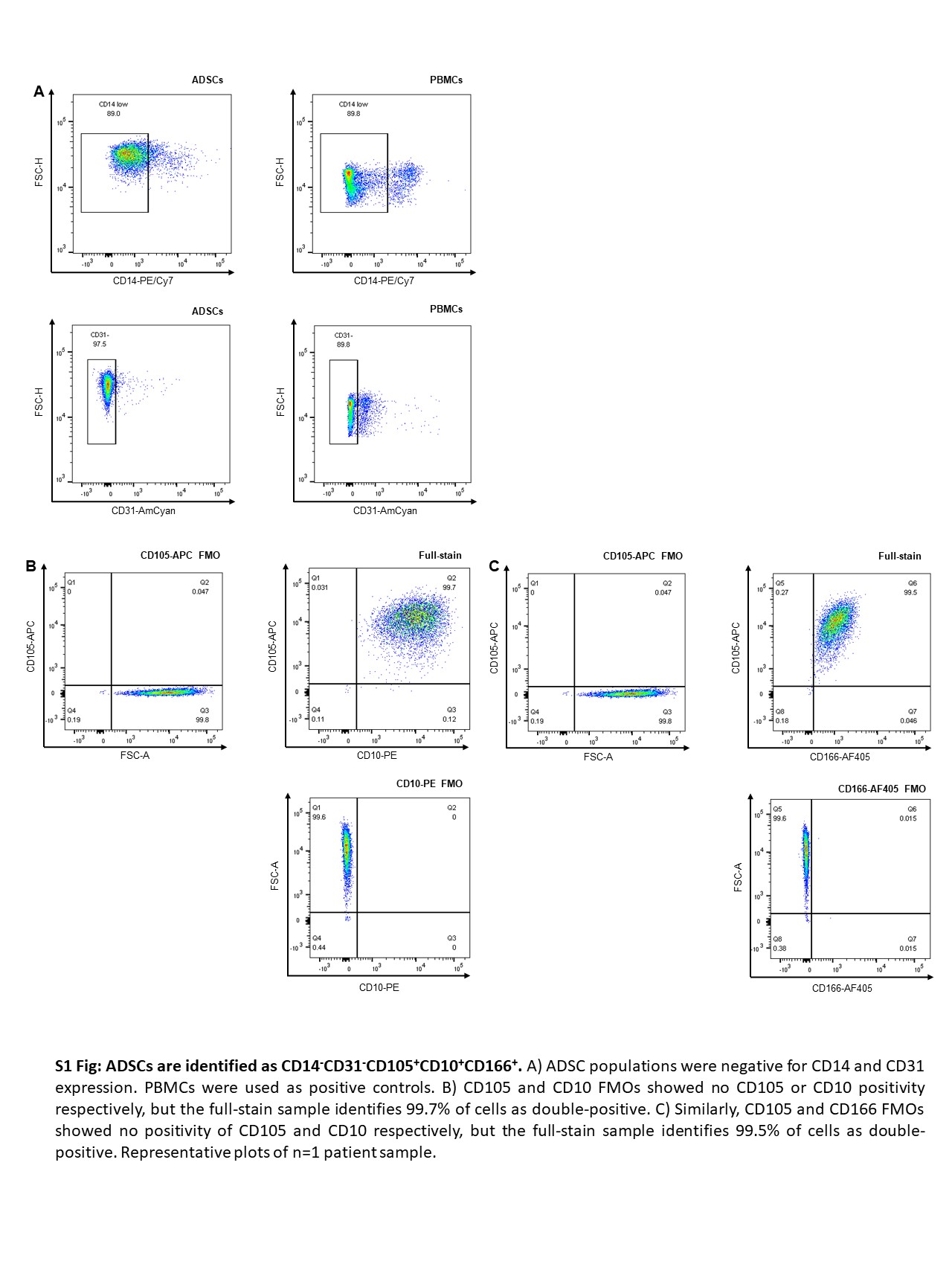

**S1 Fig: ADSCs are identified as CD14^-^CD31^-^CD105^+^CD10^+^CD166^+^. A**) ADSC populations were negative for CD14 and CD31 expression. PBMCs were used as positive controls. B) CD105 and CD10 FMOs showed no CD105 or CD10 positivity respectively, but the full-stain sample identifies 99.7% of cells as double-positive. C) Similarly, CD105 and CD166 FMOs showed no positivity of CD105 and CD10 respectively, but the full-stain sample identifies 99.5% of cells as double-positive. Representative plots of n=1 patient sample.

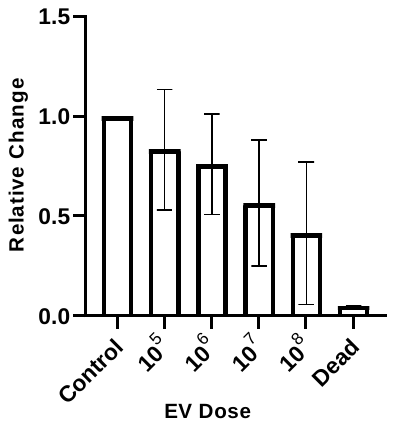

**S2 Fig: ADSC-EVs facilitate decreased macrophage cell count as dose increases.** ADSC-EVs were isolated from two AFG patients, titrated from 1x10^5^ to 1x10^8^ particles/mL and added separately to M0-like macrophage cultures from two healthy volunteers. Cell counts were assessed on a spectral cytometer (Cytek). All values are relative to a D-EV control. Heat-killed macrophages were used as a positive control (“dead”). Results are expressed as mean ± SD.

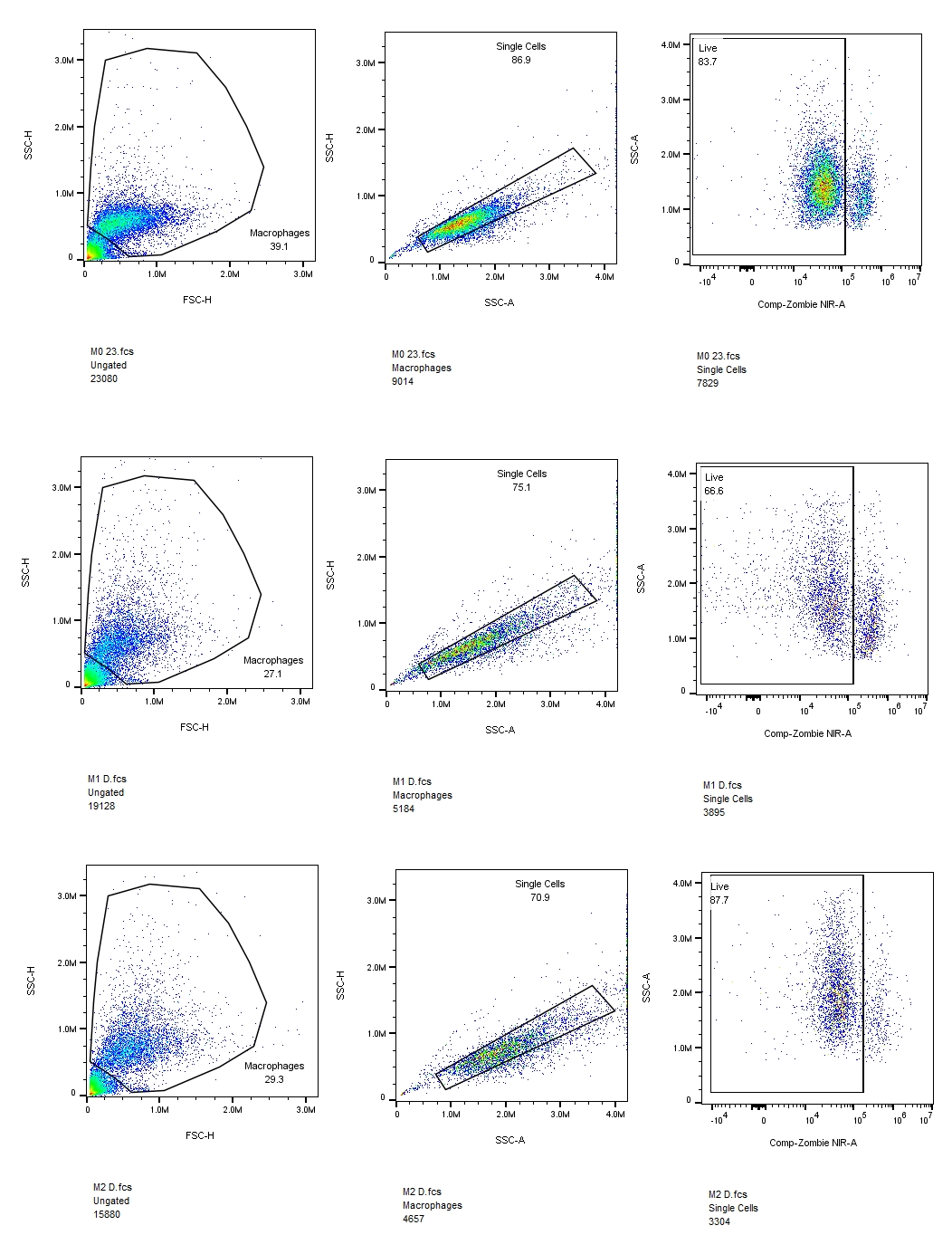

**S3 Fig: Gating strategy to identify live macrophage population.** Macrophages were identified by their side scatter (SSC-H) and forward scatter (FSC-H) properties. Doublet exclusion was performed by examining the SSC-H vs. SSC-A profile. Live cells were gated on based on their negative expression of Zombie-NIR.

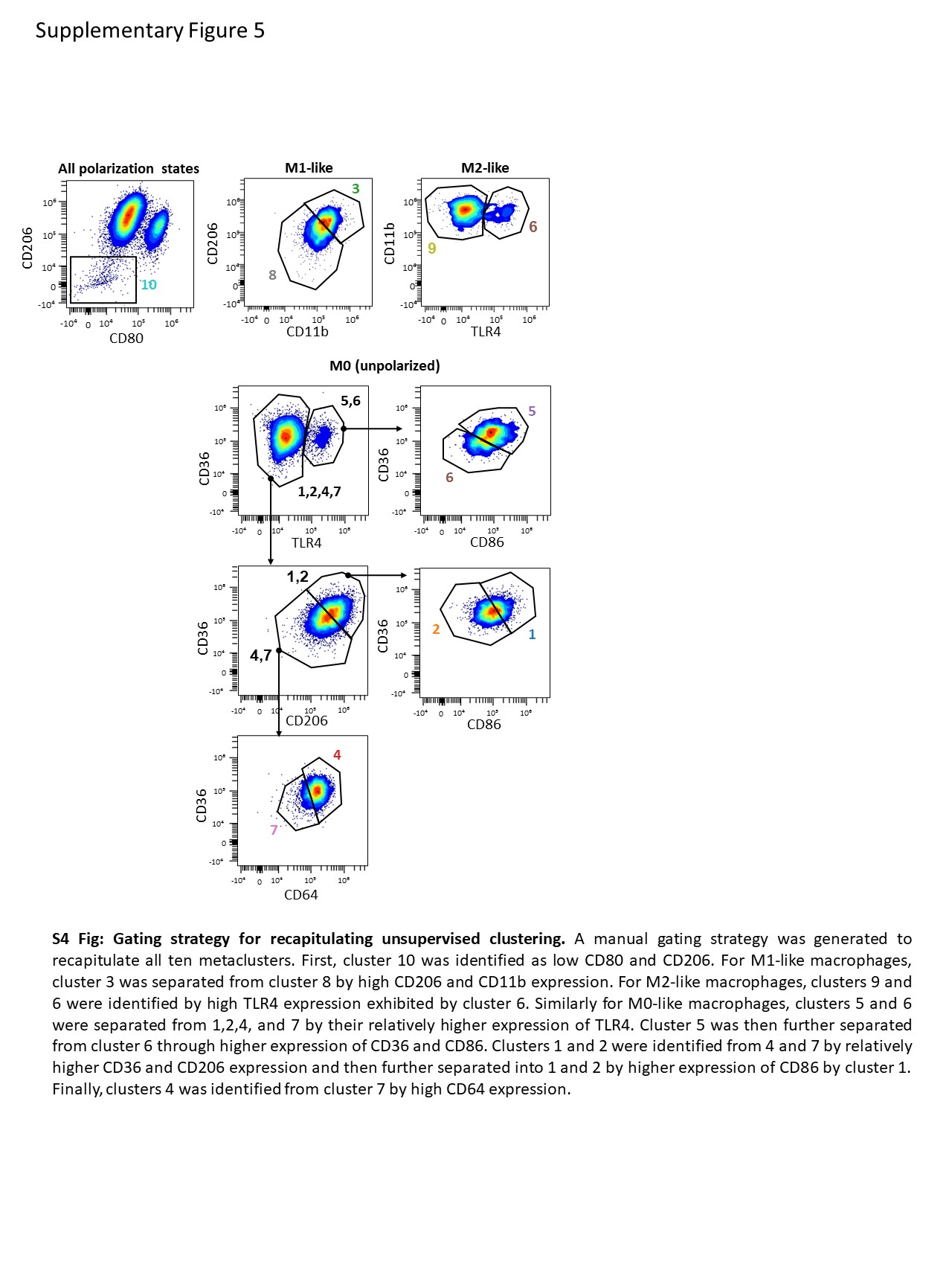

**S4 Fig: Gating strategy for recapitulating unsupervised clustering.** A manual gating strategy was generated to recapitulate all ten metaclusters. First, cluster 10 was identified as low CD80 and CD206. For M1-like macrophages, cluster 3 was separated from cluster 8 by high CD206 and CD11b expression. For M2-like macrophages, clusters 9 and 6 were identified by high TLR4 expression exhibited by cluster 6. Similarly for M0-like macrophages, clusters 5 and 6 were separated from 1,2,4, and 7 by their relatively higher expression of TLR4. Cluster 5 was then further separated from cluster 6 through higher expression of CD36 and CD86. Clusters 1 and 2 were identified from 4 and 7 by relatively higher CD36 and CD206 expression and then further separated into 1 and 2 by higher expression of CD86 by cluster 1. Finally, clusters 4 was identified from cluster 7 by high CD64 expression.

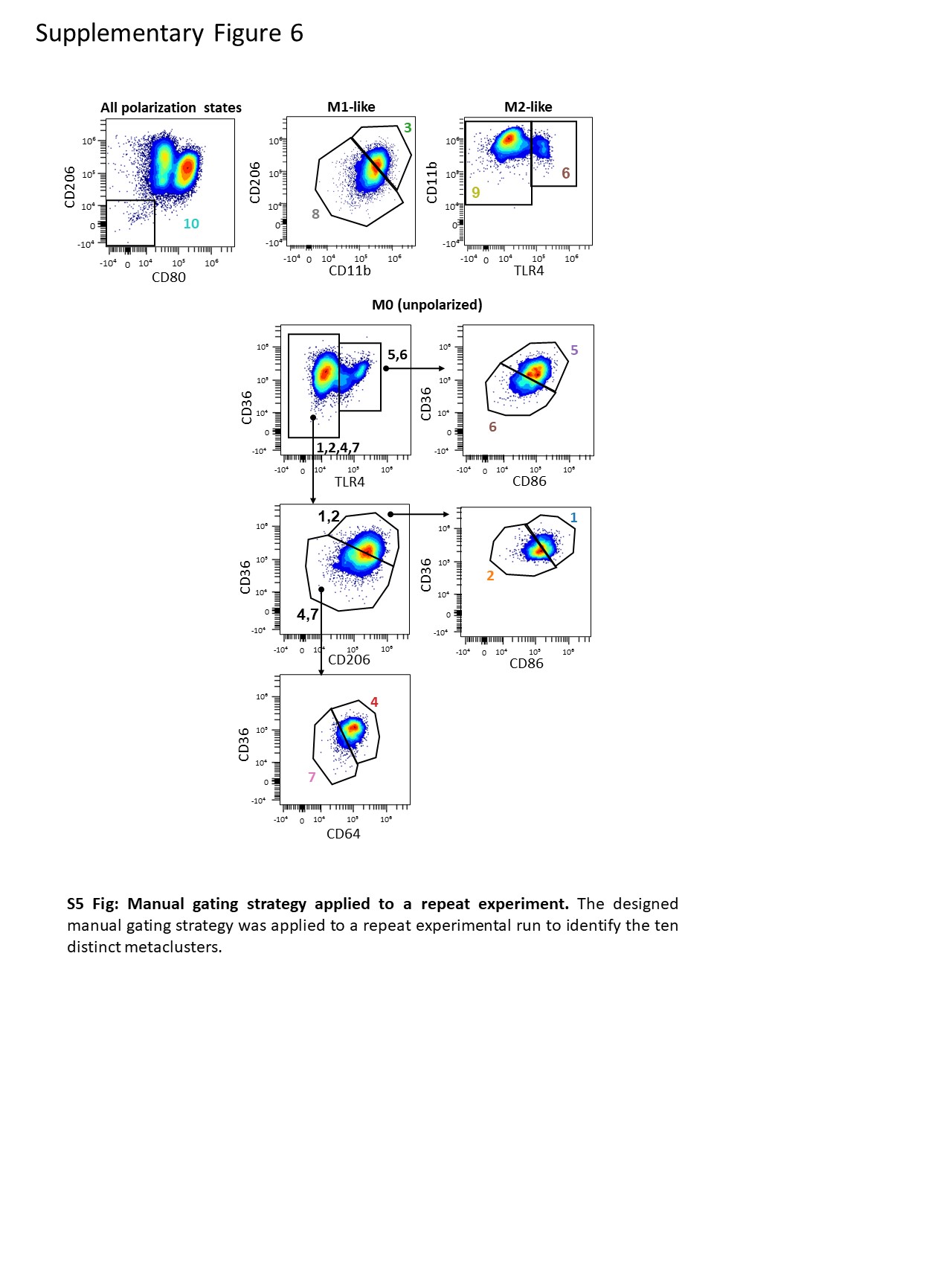

**S5 Fig: Manual gating strategy applied to a repeat experiment.** The designed manual gating strategy was applied to a repeat experimental run to identify the ten distinct metaclusters.

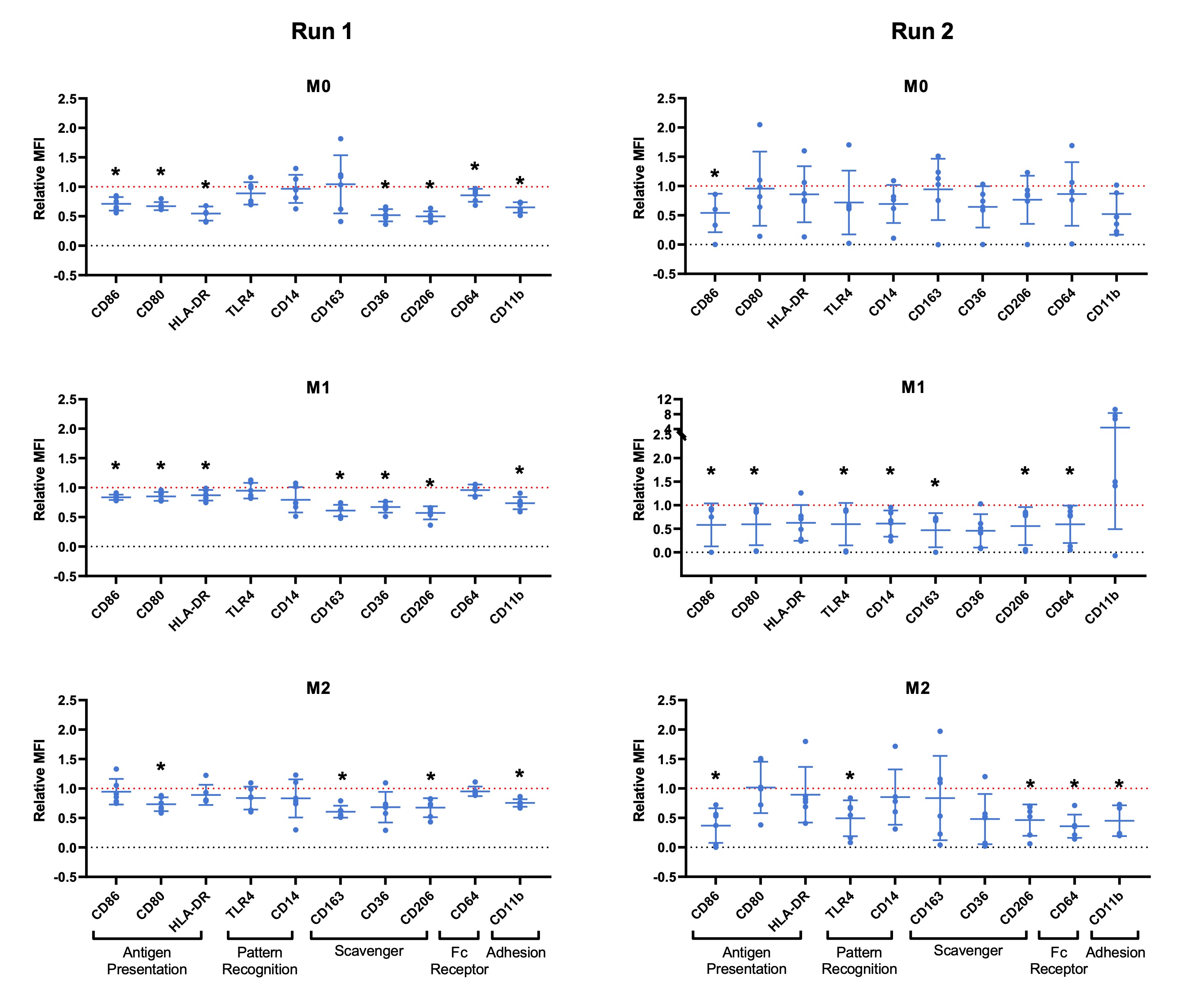

**S6 Fig: Individual marker analysis on a repeat experimental run.** When a second experimental run was analysed, ADSC-EVs reduce the expression of most individual markers relative to D-EVs (red line). Data were analysed using unpaired t tests. *p*<0.05 was deemed statistically significant.

**S1 Table: Western blot antibodies**

| **Antibody Target (clone)** | **Host/Isotype** | **Dilution in 3% BSA in T-TBS** | **Company** |
| --- | --- | --- | --- |
| CD9 | Rabbit/IgG | 1:200 | Invitrogen |
| TSG101 | Rabbit/IgG | 1:200 | Invitrogen |
| Apolipoprotein B | Rabbit/IgG | 1:200 | Invitrogen |
| Calnexin | Rabbit/IgG | 1:200 | Invitrogen |
| Goat anti-rabbit secondary antibody | Goat/IgG | 1:1000 | Invitrogen |

**S2 Table: Flow cytometry macrophage panel**

| **Marker** | **Fluorophore** | **Catalogue #** | **Company** | **Titration** | **Reason for panel incorporation** |
| --- | --- | --- | --- | --- | --- |
| CD86 | BB700 | 566474 | BD Bioscience | 1:100 | Putative M1 marker, antigen presentation, T cell activation |
| CD80 | BV510 | 557223 | BD Bioscience | 1:100 | Putative M1 marker, antigen presentation, T cell activation |
| HLA-DR | BV480 | 566154 | BD Bioscience | 1:640 | Antigen presentation and T cell activation |
| TLR4 | AF700 | 56991742 | Thermo Fisher Scientific | 1:20 | Pattern recognition |
| CD14 | BV570 | 301832 | Biolegend | 1:40 | Pattern recognition |
| CD163 | BV421 | 562643 | BD Bioscience | 1:400 | Putative M2 marker, scavenger receptor |
| CD36 | APC | 550956 | BD Bioscience | 1:2.5 | Putative M2 marker, scavenger receptor |
| CD206 | PE | 555954 | BD Bioscience | 1:20 | Putative M2 marker, scavenger receptor |
| CD64 | BV605 | 305034 | Biolegend | 1:40 | Putative M1 marker, Fc receptor |
| CD11b | BB515 | 564517 | BD Bioscience | 1:20 | Cellular adhesion |
| Zombie NIR |  | 423105 | Biolegend | 1:1000 | Live/Dead |

**S3 Table: Flow cytometry ADSC panel**

| **Antibody Target (clone)** | **Isotype** | **Laser (fluorophore)** | **Company** | **Dilution in 100 µL staining volume** | **Specificity to ADSCs?** | **Expected to be present on live ADSCs?** |
| --- | --- | --- | --- | --- | --- | --- |
| CD166 (3A6) | Mouse IgG_1_, κ | Violet (AF405) | BD Bioscience (CA, USA) | 1:20 | Marker of cells with high proliferative capabilities | Yes |
| CD105 (266) | Mouse IgG_1_, κ | Red (APC) | BD Bioscience | 1:20 | Stem cell marker | Yes |
| CD10 (HI10a) | Mouse IgG_1_, κ | Blue (PE) | BD Bioscience | 1:5 | Marker of adipogenic capabilities | Yes |
| CD31 (WM59) | Mouse IgG_1_, κ | Violet (AmCyan) | BD Bioscience | 1:20 | Endothelial cell marker | No |
| CD14 (MϕP9) | Mouse IgG_2b_, κ | Blue (PECy7) | BD Bioscience | 1:20 | Monocyte / macrophage marker | No |
| FVS780 |  | Red (APC-Cy7 ) | BD Bioscience | 1:1000 |  | No |

**Supplementary Methods**

At passage three, ADSCs from three consecutive patients were lifted using 0.25% Trypsin-EDTA, incubated at 37^◦^C for 2 mins, and centrifuged at 1000 x g for 4 mins. The supernatant was discarded and the pellet was resuspended in the desired diluted antibodies. Antibodies, outlined in Supplementary Table 3, were diluted in staining buffer (2% FBS, 98% PBS) and cells were stained at 4^◦^C for 30 mins. As outlined in Supplementary Table 1, CD10, CD105, and CD166 were used as ADSC-specific markers, while CD14 and CD31 were used to exclude contaminating cells. FVS780 was added at the same time as antibodies to prevent cell loss from multiple washes. ADSCs were fixed in 1% paraformaldehyde (PFA) at 4^◦^C for 20 mins, centrifuged at 400 x g for 12 mins, and resuspended in staining buffer. Samples were analysed on a FACSCanto II Flow Cytometer (Becton Dickinson) and analysed using FlowJo software (version 10.7.1). Fluorescent minus one (FMO) controls were used to delineate positivity and peripheral blood mononuclear cells (PBMCs) from a single healthy subject were used as a positive control for CD14 and CD31. Live, single ADSCs were identified as represented in the gating strategy in Supplementary Figure 2. PBMCs were also identified by gating on FSC vs SSC, and removing singlets and dead cells.
